# IP_3_/Ca^2+^ signals regulate larval to pupal transition under nutrient stress through the H3K36 methyltransferase dSET2

**DOI:** 10.1101/2020.11.25.399329

**Authors:** Rishav Mitra, Shlesha Richhariya, Siddharth Jayakumar, Dimple Notani, Gaiti Hasan

**Affiliations:** National Centre for Biological Sciences, Tata Institute of Fundamental Research, Bangalore, India; Department of Biology, Brandeis University, Waltham, USA; Harvard University, Cambridge, USA

**Author notes:** **Summary statement** A positive feedback loop between Ca^2+^ signaling and a histone modifier enhances gene expression in key larval neurons and drives larval to pupal transitions on a protein deficient diet.

**Keywords:** Gene expression, muscarinic acetylcholine receptor, IP_3_ receptor, neuronal epigenetics

## Abstract

Persistent loss of dietary protein usually signals a shutdown of key metabolic pathways. In *Drosophila* larvae, that have crossed “critical weight” and can pupariate to form viable adults, such a metabolic shut-down would needlessly lead to death. IP_3_/Ca^2+^ signals in certain interneurons (*vGlut^VGN634^*^1^) allow *Drosophila* larvae to pupariate on a protein-deficient diet by partially circumventing this shutdown through upregulation of neuropeptide signaling and the expression of ecdysone synthesis genes. Here we show that IP_3_/Ca^2+^ signals in *vGlut^VGN634^*^1^ neurons drive expression of *dSET2*, a *Drosophila* Histone 3 Lysine 36 methyltransferase. Further, *dSET2* expression is required for larvae to pupariate in the absence of dietary protein. IP_3_/Ca^2+^ signal-driven *dSET2* expression upregulates key Ca^2+^ signaling genes through a novel positive feedback loop. Transcriptomic studies coupled with analysis of existing ChIP-seq datasets identified genes from larval and pupal stages, that normally exhibit robust H3K36 trimethyl marks on their gene bodies and concomitantly undergo stronger downregulation by knockdown of either an intracellular Ca^2+^ release channel the IP_3_R or dSET2. IP_3_/Ca^2+^ signals thus regulate gene expression through dSET2 mediated H3K36 marks on select neuronal genes for the larval to pupal transition.

## Introduction

Integration of nutrient uptake with developmental processes is essential for organismal growth and survival (Britton et al., 2002; Wang & Lei, 2018; Ward & Thompson, 2012). In holometabolous insects such as *Drosophila* growth occurs primarily in the larval stages and is heavily dependent on access to nutrients, until the onset of pupariation which is a nutrient-independent period of development (Nijhout, 2003). Larvae that have reached the “critical weight” checkpoint can overcome protein deprivation and pupariate (Boulan et al., 2015; Mirth et al., 2005). Pupariation on a protein deficient diet requires acetylcholine stimulated Ca^2+^-release from the inositol 1,4,5-trisphosphate receptor (IP_3_R) in a subset of interneurons (Jayakumar et al., 2016). Rhythmic IP_3_/Ca^2+^ signals in the interneuron subset stimulate neuropeptide release for upregulation of ecdysone synthesis genes and interestingly also help maintain neuronal gene expression (Jayakumar et al., 2016, 2018).

Gene expression changes in neurons are primarily thought to occur by activity driven transcription of immediate-early genes and their downstream signaling mechanisms (Chen et al., 2016; Dolmetsch, 2003). Little notice has been given to how IP_3_/Ca^2+^ changes, downstream of myriad signaling pathways initiated by metabotropic receptors, regulate neuronal gene expression. We report here an essential role for the Histone 3 Lys 36 methyltransferase (H3K36me), dSET2 (Fig 1A), downstream of neuronal IP_3_/Ca^2+^ signals, for pupariation on a protein deficient diet. In *Drosophila*, as well as in other organisms (McDaniel et al., 2017), H3K36 trimethylation by dSET2 (or its orthologues; Suppl Fig 1A, B) is required for efficient transcriptional elongation (Schaft et al., 2003) and serves to upregulate gene expression (Bannister et al., 2005).

**Figure 1:**
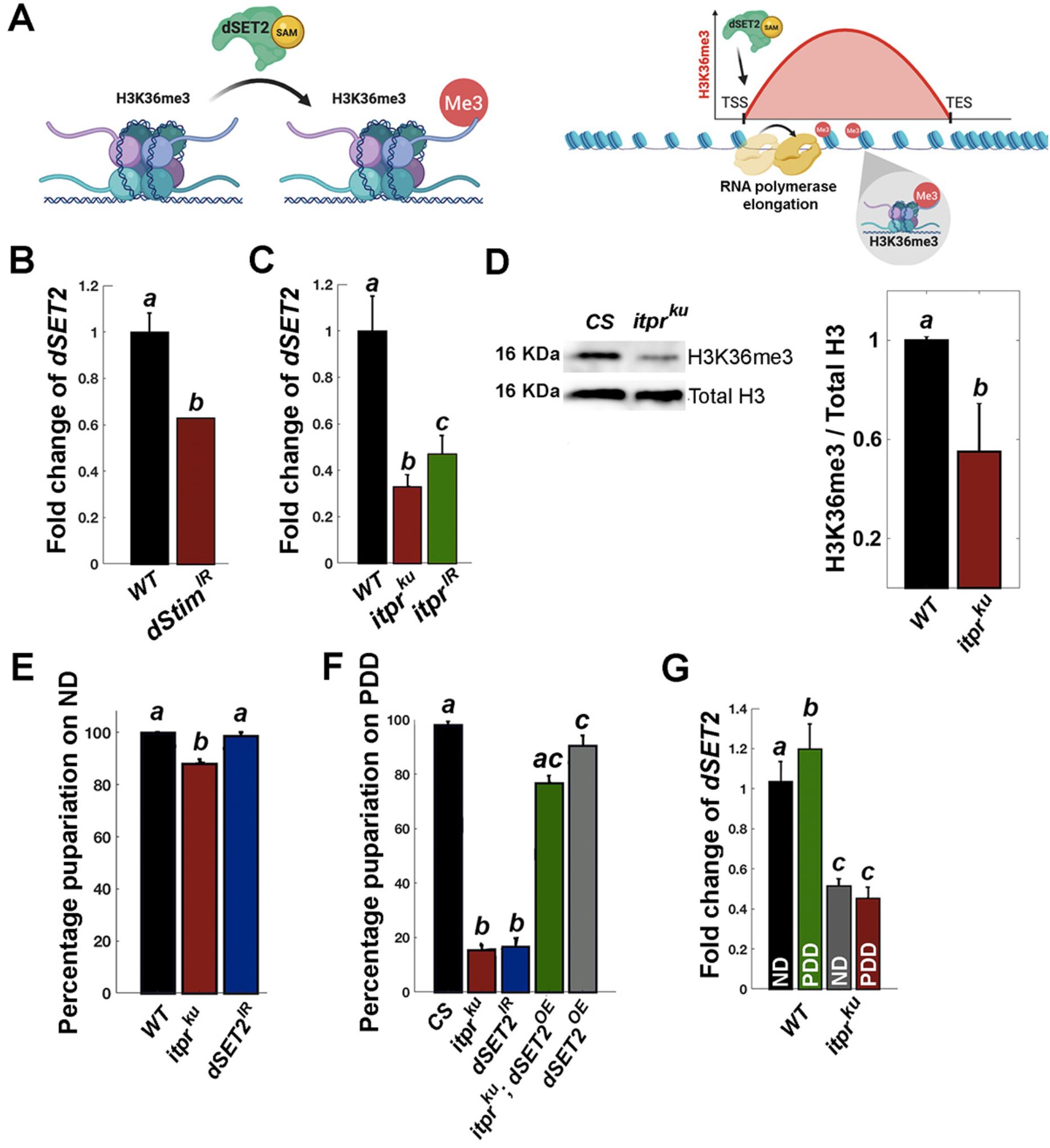
The IP_3_R is required for neuronal expression of dSET2. A) dSET2 is a H3K36 methyltransferase (left). Enrichment of trimethylated H3K36 occurs along the gene body is implicated in transcriptional elongation (Schaft et al., 2003) (right). B) *dSET2* levels are significantly attenuated upon RNAi-driven knockdown of the ER-Ca^2+^ sensor *dSTIM* (*dSTIM^IR^*) in *Drosophila* pupal brains as compared to the control genotype C) *dSET2* levels also go down significantly in larval brains with reduced IP_3_R function obtained from the viable IP_3_R mutant combination, *itpr^ka1091/ug3^* (or *itpr^ku^*) and pan-neuronal knockdown of the IP_3_R (*itpr^IR^*). D) Histone 3 Lysine 36 trimethylation (H3K36me3) is significantly reduced in larval brain lysates from *itpr^ku^* as evident in a representative western blot (left) and quantified on the right. E) Pupariation on a normal diet (ND) is affected mildly in *itpr^ku^* and unaffected by knockdown of *dSET2* in *vGlut^VGN6341^*neurons. F) Pupariation on a protein deprived diet (PDD) is greatly reduced in *itpr^ku^* animals and by knockdown of *dSET2* in *vGlut^VGN6341^* neurons. Overexpression of *dSET2* in *vGlut^VGN6341^*neurons rescues pupariation of *itpr^ku^* larvae. G) Expression of *dSET2* is not affected by dietary regimes. All bar graphs represent mean values (+SEM) from a minimum of three experiments. The same letter above the bars represents statistically indistinguishable groups after performing ANOVA and a post-hoc Tukey test (p<0.05).

## Results and Discussion

### IP_3_R regulates neuronal expression of dSET2 a Histone 3 Lysine 36 methyltransferase

Regulation of *dSET2* expression, by intracellular Ca^2+^ signaling was first suggested in a transcriptomic screen from the *Drosophila* pupal central nervous system with pan-neuronal knockdown of the Endoplasmic Reticulum (ER) Ca^2+^ sensor *dSTIM* (Fig 1B; Richhariya et al., 2017). To test if *dSET2* expression might be altered by Ca^2+^ release from the ER, we measured levels of *dSET2* mRNA in IP_3_R mutant *Drosophila* larvae (*itpr^ka1091/ug3^* or *itpr^ku^*; Joshi et al., 2004), known to exhibit reduced IP_3_-mediated Ca^2+^ release upon stimulation of the muscarinic acetylcholine receptor or mAchR (Jayakumar et al., 2016). Larval *itpr^ku^* brains exhibit greater than 50% reduction in levels of *dSET2* mRNA (Fig 1C). A similar decrease in *dSET2* levels was observed in larval brains with pan-neuronal knock down of the IP_3_R (*elav^C155^GAL4>itpr^IR^*; Fig 1C) suggesting that *dSET2* expression in neurons is indeed sensitive to IP_3_-mediated Ca^2+^ release through the IP_3_R.

The *Drosophila* genome encodes a single H3K36 trimethylation (H3K36me3) protein, *dSET2* (Fig S1A, B; Stabell et al., 2007). Hence, the extent of H3K36me3 is a direct measure of dSET2 activity. We measured levels of H3K36me3 in lysates from *itpr^ku^* mutant larval brains where quantitative measurement revealed a significant reduction in trimethylated H3K36 (Fig 1D). Thus, reduced IP_3_R function in *Drosophila* larvae resulted in reduced amounts of *dSET2* mRNA, accompanied by hypomethylated H3K36 indicating reduction in dSET2 protein. To study dSET2 function in the larval brain, we used two independent RNAi lines, *dSET2^IR-1^* and *dSET2^IR-2^*. Pan-neuronal knockdowns for each resulted in significant down-regulation of *dSET2* in the brain, whereas overexpression of dSET2 (*dSET2^OE^*) achieved a 1.8-fold increase in expression (Fig S1C).

From previous studies we know that IP_3_-mediated Ca^2+^ release enables larval-pupal transitions under protein deprivation and the focus of this deficit lies in a subset of glutamatergic interneurons, *vGlut^VGN6341^,* located in the ventral ganglion (Jayakumar et al., 2016; 2018). To assess the physiological consequences of deficient H3K36 trimethylation, we measured pupariation on normal (ND) and protein deficient diets (PDD). All wild type (CS) larvae on ND (Fig 1E) and PDD (Fig 1F) when transferred as mid-third instars, progress through development to form pupae. Specific depletion of *dSET2* in *vGlut^VGN6341^* neurons by expression of an RNAi (*dSET2^IR-2^*) has no effect on pupariation rates on ND, and in agreement with earlier findings *itpr^ku^* mutant larvae show a minor decrease in pupariation on ND (Fig 1E). However, when mid-third instar larvae with knockdown of *dSET2* in *vGlut^VGN6341^* neurons, were switched from ND to PDD, there was a significant reduction of pupariation (Fig 1F) as observed previously for *itpr^ku^* larvae (Jayakumar et al., 2016). Remarkably, reduced pupariation on PDD in *itpr^ku^* larvae could be rescued from <20% to >70% by overexpressing *dSET2* in *vGlut^VGN634^*^1^ neurons (Fig 1F). Thus, a systemic deficit due to loss of intracellular Ca^2+^ signaling in IP_3_R mutant larvae can be compensated by overexpression of dSET2 in a subset of glutamatergic interneurons.

*dSET2* levels in the brains of wild type larvae, either on PDD or ND, were similar (Fig 1G). In *itpr^ku^* mutant brains, however, *dSET2* mRNA levels are significantly down-regulated in both dietary regimes (Fig 1G), indicating that attenuated ER Ca^2+^ release through the IP_3_R, affects the expression of *dSET2,* independent of diet.

### Ca^2+^ release through the IP_3_ receptor is attenuated in neurons with reduced dSET2

Peripheral cholinergic inputs signal the loss of dietary protein and stimulate IP_3_/Ca^2+^ signaling through the mAchR in *vGlut^VGN6341^* neurons (Fig 2A; Jayakumar et al., 2016). The cellular consequence of reduced dSET2 in *vGlut^VGN634^*^1^ neurons was examined next by calcium imaging in ex-vivo larval brains (Fig 2B; Fig S2). Calcium responses were recorded upon stimulation with 50 μM carbachol (CCh), an agonist for the mAchR, from *vGlut^VGN634^*^1^ interneurons using GCaMP6m (Chen et al., 2013) (Fig 2C). *vGlut^VGN634^*^1^ neurons from wild-type larvae exhibit a robust calcium transient upon carbachol stimulation (Jayakumar et al., 2016) (Fig 2C-E). Knockdown of either *dSET2* (Fig 2C, 2D) or the IP_3_R (Fig 2E) in *vGlut^VGN634^*^1^ neurons, attenuated the calcium response to carbachol to a significant extent, as evident from reduced peak values and areas under the curve, in comparison with WT controls (Fig 2F, G). These data demonstrate that loss of dSET2 in *vGlut^VGN6341^* neurons impedes Ca^2+^ release through the IP_3_R in response to carbachol stimulation.

**Figure 2:**
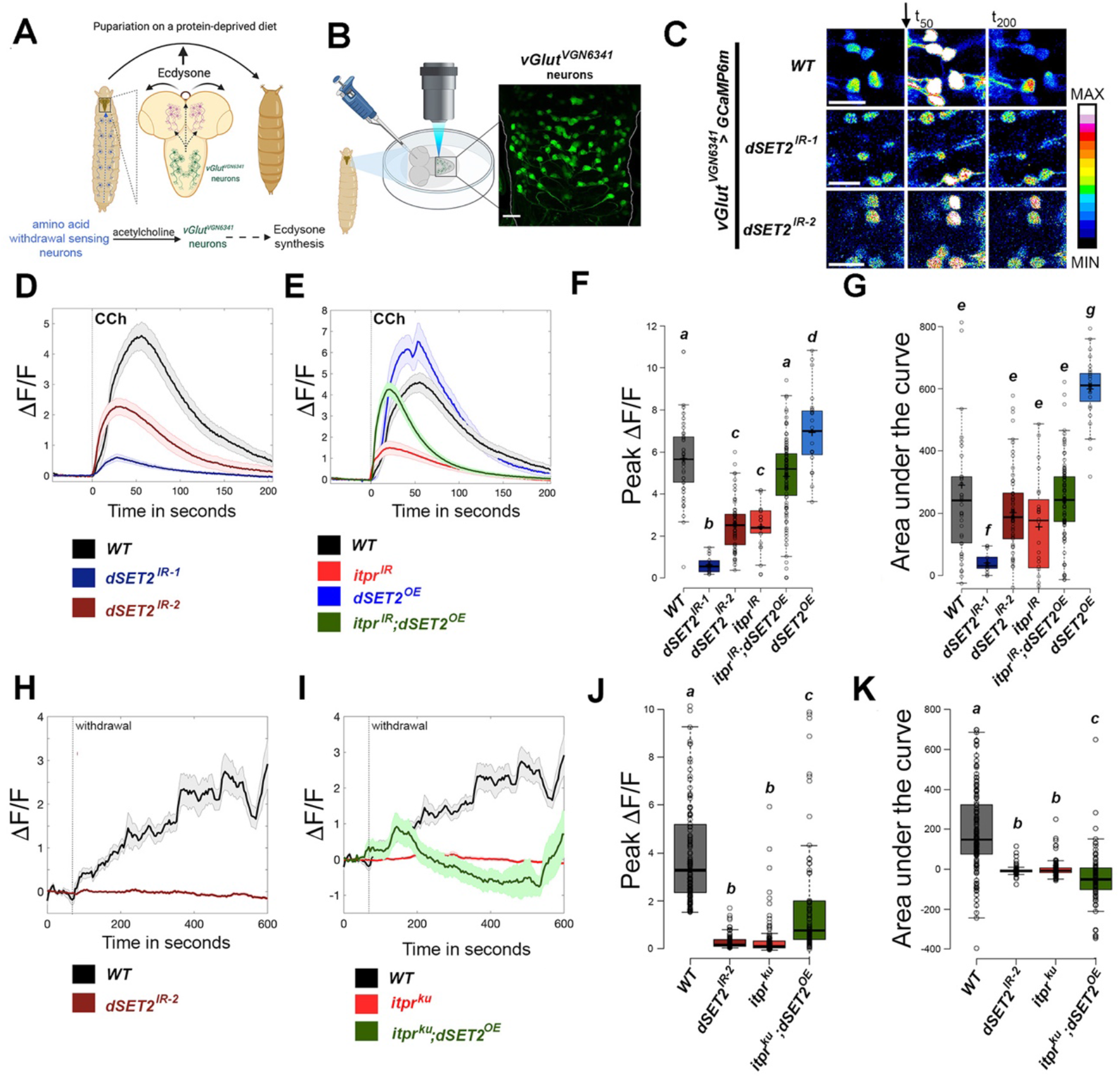
IP_3_ mediated Ca^2+^ responses in *vGlut^VGN6341^* neurons require dSET2. A) *vGlut^VGN6341^* neurons mediate larval to pupal transitions of protein deprived larvae that have gone past critical weight. B) Experimental setup for ex-vivo imaging from larval *vGlut^VGN6341^* neurons. C) Knockdown of *dSET2* dampens carbachol (50μM) evoked calcium transients from *vGlut^VGN6341^* cells. D, E) Graphs depict the average response to 50 μM carbachol (CCh) of *vGlut^VGN634^* neurons from the indicated genotypes, visualized by changes in GCamP6m fluorescence (ΔF/F) over time. *dSET2^OE^* indicates overexpression of *dSET2.* F) Peak responses to carbachol and G) Areas under the curve for *vGlut^VGN6341^* neurons of the indicated genotypes. H, I) Graphs depict the average response of *vGlut^VGN6341^* neurons to withdrawal of ambient amino acids in the indicated genotypes. J) Peak responses to carbachol and K) Areas under the curve for *vGlut^VGN6341^* neurons of the indicated genotypes. Boxes indicate the 25th and 75th percentiles and whiskers extend to 1.5 times the interquartile range. Lines in the center show the median. Different letters indicate statistically significant differences (p<0.05) after performing ANOVA and a post hoc Tukey test.

Based on rescue of pupariation of IP_3_R mutants by dSET2 (Fig 1F), next we tested Ca^2+^ responses upon over-expression of *dSET2* in *vGlut^VGN6341^* neurons. Indeed, the attenuated Ca^2+^ response seen in *vGlut^VGN6341^* neurons with *itpr^IR^*, post-carbachol stimulation, was rescued back to wild type levels by overexpression of *dSET2* (Fig 2D), as evident from the peak values and area under the curve (Fig 2F; Fig S2). Over-expression of *dSET2* in wild type *vGlut^VGN6341^* neurons resulted in significant increase in the evoked response to carbachol as compared to WT (Fig 2E-G). Acute amino acid withdrawal evokes a series of slow calcium transients in *vGlut^VGN6341^* that are lost in *itpr^ku^* (Fig 2H). Initiation of these Ca^2+^ transients requires cholinergic inputs whereas long term propagation depends upon neuropeptide receptors (Jayakumar et al., 2018). Knockdown of *dSET2* (*dSET2^IR-2^)* abrogated the Ca^2+^ transients whereas overexpression of *dSET2* in *vGlut^VGN6341^* neurons of *itpr^ku^* restored the initial, but not the longer, response. (Fig 2H-K). These data demonstrate that dSET2 rescues the cholinergic response in *vGlut^VGN6341^* neurons.

### dSET2 regulates expression of the muscarinic acetylcholine receptor and the IP_3_R in *vGlut^VGN634^*^1^ neurons

Expression of *dSET2* is regulated by IP_3_/Ca^2+^ signaling (summarized in Fig 3A). However, loss of carbachol-stimulated IP_3_/Ca^2+^ signals upon knockdown of *dSET2* followed by their rescue upon overexpression of *dSET2,* suggests feedback regulation by dSET2 of key components of IP_3_/Ca^2+^ signaling in neurons. This idea was investigated next by measuring transcript levels of *mAchR* and *itpr*, two critical components of the pathway. Brains with pan-neuronal knock-down of *dSET2* did not exhibit a detectable change in the expression of either *mAchR* or *itpr* (Fig 3B). As shown previously, pan-neuronal knock-down of *itpr* resulted in a significant decrease in *dSET2* levels in the whole brain (Fig 3C). However, pan-neuronal *itpr* knockdown did not affect the expression of *mAchR* (Fig 3C).

**Figure 3.**
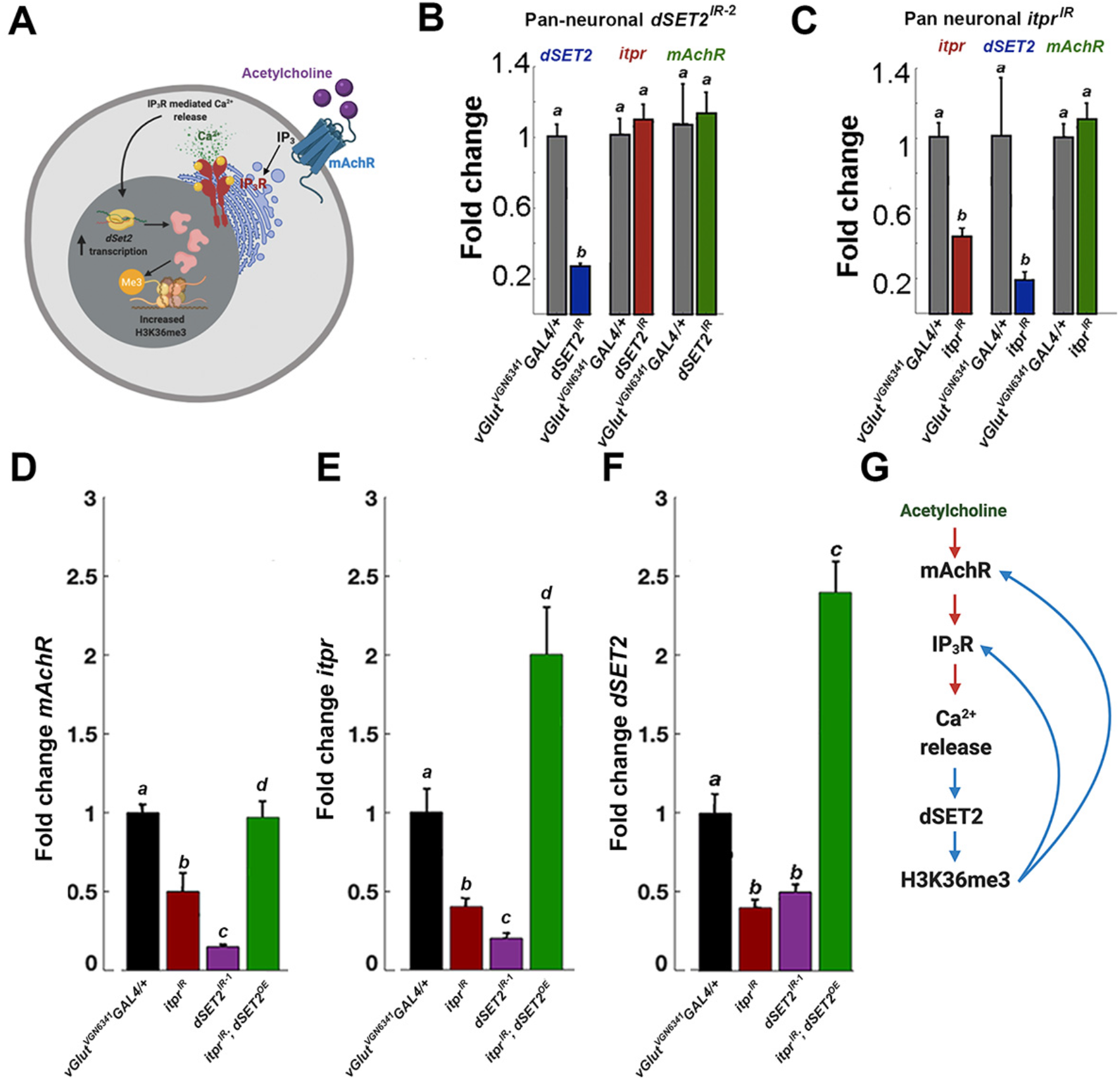
A positive feedback loop maintains the expression of dSET2, the muscarinic acetylcholine receptor and the IP_3_R in *vGlut^VGN634^*^1^ neurons. A) Model depicting transcription of *dSET2* and dSET2-mediated H3K36 trimethylation (H3K36me3) downstream of acetylcholine and IP_3_R-mediated Ca^2+^ release. B) Pan-neuronal knockdown of *dSET2* has no discernible effect on expression of *itpr* and *mAChR* mRNA in the whole brain C) Similarly, *itpr* knockdown has no effect on *mAchR* levels in the whole brain. Grey bars represent WT brains. Transcripts of *mAchR* (D), *itpr* (E) and *dSET2* (F) are downregulated in *vGlut^VGN6341^* neurons upon expression of *dSET2^IR^* (purple), *itpr^IR^* (maroon), and restored upon over-expression *of dSET2* with *itpr^IR^* (green). Black bars represent WT *vGlut^VGN6341^* neurons G) Schematic representation of the signaling pathway initiated by Ca^2+^ signals downstream of mAchR and IP_3_R that leads to elevated *dSET2* transcripts and H3K36me3. A feedback regulatory loop from dSET2 maintains expression of *mAchR* and *itpr.* Average (±SEM) fold changes of gene expression shown in the bar graphs were quantified by a minimum of three qPCR experiments. Letters above the bars represent statistically indistinguishable groups (p<0.05) after performing ANOVA and a post-hoc Tukey test.

Because both the systemic (Fig 1) and cellular phenotypes of IP_3_R and dSET2 (Fig 2) derived specifically from *vGlut^VGN634^*^1^ neurons, we hypothesized that dSET2 might regulate *mAchR* and *itpr* expression in that specific neuronal subset. In order to measure gene expression in *vGlut^VGN634^*^1^ neurons they were tagged with cytosolic eGFP and enriched by Fluorescence Activated Cell Sorting (FACS; Fig S3 A-C). This allowed for a greater than 50 - fold enrichment of GFP-tagged *vGlut^VGN634^*^1^ neurons from all required genotypes (Fig S3D).

In agreement with attenuated Ca^2+^ responses seen upon *dSET2* knockdown in *vGlut^VGN634^*^1^ neurons (Fig 2D) expression of both *mAchR* (Fig 3D) and *itpr* (Fig 3E) was downregulated by 0.25 and 0.3-fold respectively as compared with WT in sorted *vGlut^VGN634^*^1^ neurons with *dSET2* knockdown (Fig 3D, E, purple bars). These results are in contrast to pan-neuronal knockdown of *dSET2* where no change in expression of either *mAchR* or *itpr* was observed (compare expression in Fig 3B with Figs 3 E, F; purple bars). The *dSET2* RNAi (*dSET2^IR-2^*) used for these experiments reduced expression of *dSET2* (~50%) (Fig 3C) to an extent similar to pan-neuronal knockdown (Fig S1C). Moreover, *itpr* knockdown in *vGlut^VGN634^*^1^ neurons (Fig 3D, maroon) reduced expression of *mAchR* to ~ 0.5-fold, unlike pan-neuronal *itpr* knockdown where *mAchR* expression remained similar to WT (Fig 3C). The reciprocal effects of *dSET2* and *itpr* on expression of their respective genes and *mAchR*, suggests a positive feedback regulatory loop between them (Fig 3G). This idea is further supported by overexpression of *dSET2* in *vGlut^VGN634^*^1^ neurons in the presence of *itpr* RNAi (Fig 3D-F, green bars). *dSET2* expression negated the effect of the RNAi as seen by the significantly heightened expression of both *mAchR* (Fig 3D; green) and *itpr* (Fig 3E; green). These results indicate that *dSET2* and H3K36 trimethylation regulate gene expression specifically in *vGlut^VGN634^*^1^ neurons, essential for maintaining responsiveness to acetylcholine through mAchR and the IP_3_R (schematized in Fig 3G).

### *dSET2* and IP_3_/Ca^2+^ signals together regulate a subset of the neuronal transcriptome

To understand the breadth of neuronal gene expression changes mediated by IP_3_/Ca^2+^ and dSET2, we performed RNA-Seqs from third instar larval central nervous systems (CNS) expressing pan-neuronal *dSET2* RNAi (Fig 4A), followed by comparison with a published RNA-seq for pan-neuronal *itpr RNAi* from an equivalent stage (Fig 4B)(Jayakumar et al., 2018). *dSET2^IR^* and *itpr^IR^* conditions both exhibit greater proportion of downregulated rather than upregulated genes (Fig 4A, B, Fig S4A). Moreover, higher fold change values were observed in a greater number of downregulated as compared to upregulated genes (Fig S4A). Consequently, further analysis was performed primarily on downregulated genes. Interestingly, dysregulated genes with greater fold changes in either *dSET2 RNAi* or *itpr RNAi* conditions exhibited higher baseline expression as measured from their respective control genotypes (*RNAi/+*; Fig S4B) suggesting that appropriate regulation of developmental gene expression in the larval brain includes components of IP_3_R/Ca^2+^ signaling and dSET2 mediated H3K36me3 modification.

**Figure 4.**
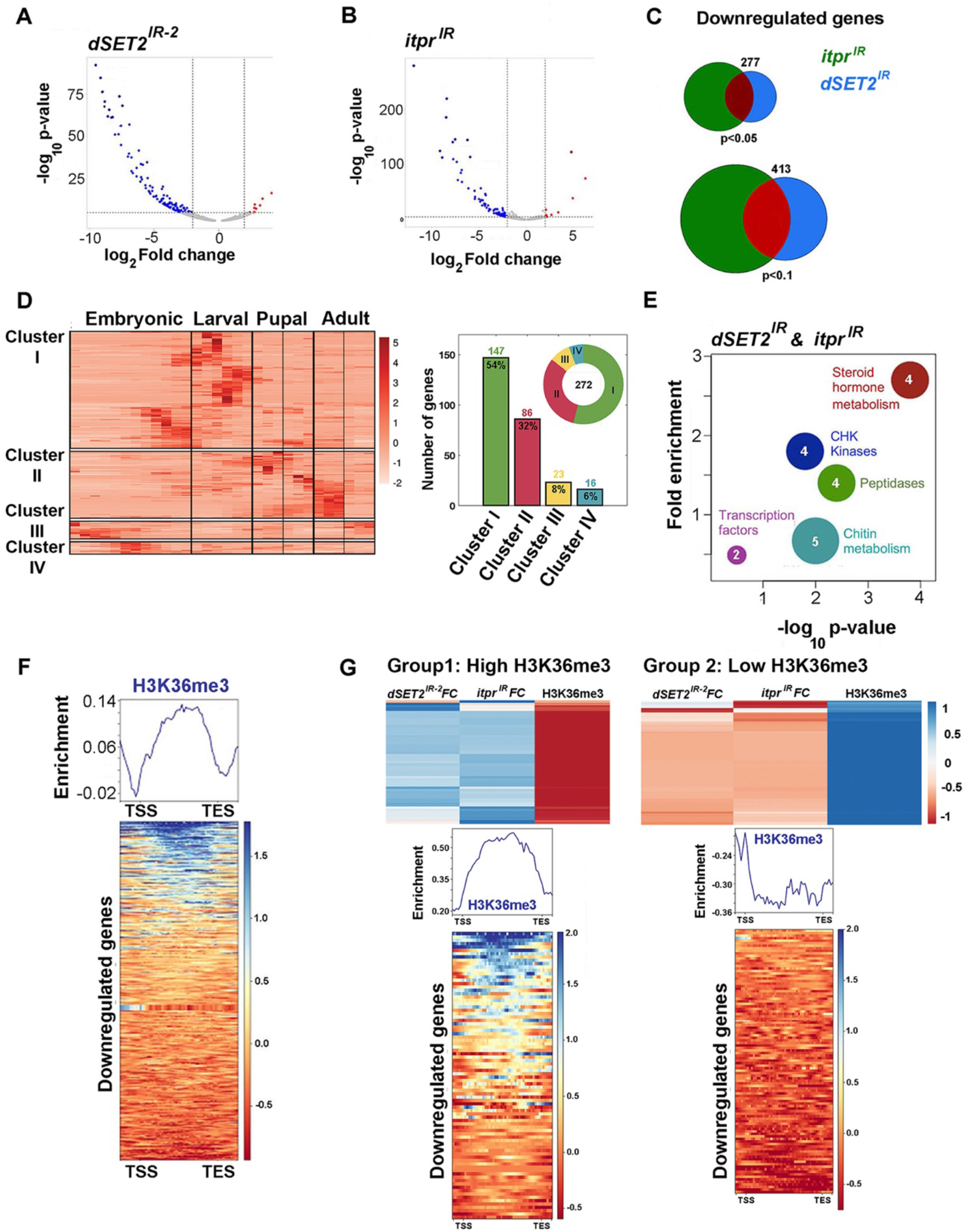
*dSET2* and IP_3_/Ca^2+^ signals together regulate a subset of the neuronal transcriptome. Differentially expressed (DE) genes in the larval CNS upon A) *dSET2* knockdown and *itpr* knockdown are primarily down-regulated. C) A common set of down-regulated genes identified upon knockdown of *dSET2 and itpr.* D) A majority of genes down-regulated in common between *dSET2^IR^* and *itpr^IR^* conditions show peak baseline expression in larvae and pupae. E) Gene Ontology (GO) analysis of genes down-regulated in common between *itpr^IR^* and *dSET2^IR^* conditions. Number of genes for each GO term is shown within the circle. F) Relative enrichment of H3K36me3 signal is evident across the gene body in the *Drosophila* BG3 cell line (Celniker et al., 2009; see methods) among genes down-regulated in common between *itpr^IR^* and *dSET2^IR^* conditions. G) Greater enrichment of H3K36me3 is found in DE genes with higher fold change values (FC) from *dSET2^IR-2^* and *itpr^IR^* RNA seqs (left). DE genes with smaller FCs correlate with reduced H3K36me3 marks. The scale indicates difference between the respective fold changes and H3K36me3 enrichment (in DmBGL3 cells; from Karpen et al., 2009) over the gene body, with unit variance scaling.

Next, we compared down-regulated genes from *dSET2^IR^* and *itpr^IR^* datasets and identified 277 genes in common (Fig 4C; p<0.05). At a lower stringency (p<0.1) the number of common genes increased to 413. A developmental expression profile of the 277 down-regulated genes (p<0.05) in wild-type *Drosophila* (Celniker et al., 2009) identified 4 clusters with differential peaks of expression (Fig 4D). Among these, a large number (233 of 277 or 86%; clusters I and II) also express in larvae and pupae, re-iterating the likely importance of IP_3_/Ca^2+^ signals and H3K36 trimethylation during these developmental stages.

Gene ontology and pathway analysis of the 277 common downregulated genes revealed a significant enrichment in GO categories that are likely to aid the larval to pupal transition (Fig 4E) such as ecdysone synthesis (steroid hormone metabolism), breakdown of the larval cuticle (peptidases) and synthesis of the pupal/adult cuticle (chitin metabolism). Two other GO categories of DNA damage dependent checkpoint activation (CHK Kinases) and transcription factors possibly relate to the extensive changes in gene expression profiles that occur between larvae, pupae and adults. Similar GO categories, in addition to a few others, were enriched among downregulated genes in *dSET2^IR-2^* (Fig S4C) and *itpr^IR^* (Fig S4D) datasets. Taken together these analyses of RNA-seq data support the idea that IP_3_/Ca^2+^ signals and *dSET2* expression allows for timely expression of multiple neuronal genes during the larval to pupal transition.

Based on existing knowledge that dSET2 activity catalyzes H3K36 tri-methylation on the gene body and is frequently associated with transcriptional activation and elongation (Kizer et al., 2005; Schaft et al., 2003), we hypothesized that gene bodies in common between *dSET2^IR^* and the *itpr^IR^* datasets should normally appear enriched in H3K36me3 marks. This was assessed using a published dataset from ChIP-on-chip performed in *Drosophila* ML-BG3 cells, that is derived from the 3^rd^ instar larval CNS (Celniker et al., 2009). For this analysis, we selected commonly downregulated genes (413; p<0.1) identified from *dSET2* and *itpr* knockdown datasets (Fig 4C). An appreciable enrichment of H3K36me3 was observed across the gene body of these genes and as expected (Kharchenko et al., 2011; Kouzarides, 2007), we observed a drop in the flanking regions that constitute the transcriptional start and stop sites (Fig 4F).

Next, we asked if the severity of transcription loss upon *dSET2 KD* is directly proportional to the enrichment of H3K36me3 on these genes. Interestingly, such correlation yielded two categories of genes. Approximately half the genes belong to Group 1 where higher extents of downregulation in *itpr^IR^* and *dSET2^IR-2^* conditions, associate with enhanced enrichment of H3K36me3 along the gene body. These genes are likely to be direct targets of the catalytic activity of dSET2, downstream of IP_3_/Ca^2+^ signals. Group 2 represents genes with smaller fold changes and concomitant lower enrichment of H3K36me3, indicating an indirect effect of the IP_3_/Ca^2+^ and dSET2/H3K36me3 driven transcriptional program (Fig 4G).

IP_3_/Ca^2+^ signals can thus regulate changes in neuronal gene expression, through changes in histone methylation. The IP_3_R/Ca^2+^-H3K36me3 axis identified here, suggests a possible mechanism by which neurons establish a brain-wide transcriptional profile, which in *Drosophila* larvae culminates in endocrine outputs such as ecdysone biosynthesis essential for pupariation. In a broader context several studies suggest a link between intracellular Ca^2+^ signaling and histone modifications that regulate transcription (Sharma et al., 2014; Whitlock et al., 1983). Dysfunction of the IP_3_R has been implicated in certain neurodegenerative conditions such as Alzheimer’s, Huntington’s disease and Spino-cerebellar ataxias (Egorova & Bezprozvanny, 2018; Hasan & Sharma, 2020). Rodent models for Alzheimer’s disease and accelerated senescence also show a decrease in brain wide levels of H3K36 methylation (Wang et al., 2010). Post-mortem brains obtained from Huntington’s disease patients show association of SETD2, the mammalian ortholog of dSET2, with morphological deposits linked to pathology (Passani et al., 2000) and human mutations in SETD2 have been associated with intellectual disabilities and autism spectrum disorders (Iossifov et al., 2014; Lumish et al., 2015). Future studies that investigate the IP_3_/Ca^2+^-H3K36me3 link identified here, in the context of mammalian neurons could lead to a better understanding of neurodegenerative and neurological disease pathologies and help identify potential diagnostic markers and treatment.

## Materials and Methods

### Fly strains

*Drosophila* strains were grown on standard cornmeal medium supplemented with yeast (ND) at 25°C under a light-dark cycle of 12h and 12h. The protein-deprived diet (PDD) consists of 100 mM sucrose with 1% agar. Canton S was used as wild-type (WT) throughout. All fly stocks used have been listed as a table in Supplementary Table 1.

### Generation of transgenic *UAS-dSET2*

A full-length cDNA Gold clone, LD27386, encoding dSET2 (Transcript B) was obtained from the *Drosophila* Genomic Resource Center (DGRC). The cDNA sequence was PCR amplified using Phusion enzyme (NEB, M0530S) and appropriate primers with overhangs for Not1 and Kpn1 restriction enzymes (sequences given in Supplementary Table 2). The PCR product and *pUAST-attB* vector were digested with NotI and Kpn1 restriction enzymes (NEB) and ligated to obtain the desired clones. The complete cDNA sequence was verified by Sanger sequencing. dSET2 flies were obtained by injection of embryos employing standard methods for *Drosophila* transgenics by the Fly facility at NCBS, Bengaluru.

### Pupariation assays

Larvae at 84 ± 4h post egg laying were transferred to PDD or ND in batches of 25 and were scored for pupariation. Data for each genotype on each media were obtained from at least six independent batches of larvae. These are reported as percentage pupariation.

### RNA isolation and quantitative PCR

Central nervous systems (CNS) from larvae of the appropriate genotype and age were dissected in phosphate buffer saline (PBS, 137 mM NaCl,2.7 mM KCl,10 mM Na_2_HPO_4_, 1.8 mM KH_2_PO_4_) prepared in double distilled water treated with DEPC. Each sample consisted of 5 CNSs homogenized in 500 μl of TRIzol (Ambion, ThermoFisher Scientific) per sample. At least three biological replicate samples were made for each genotype. After homogenization the sample was kept on ice and either processed further within 30 min or stored at −80°C for up to 4 weeks before processing. RNA was isolated following the manufacturer’s protocol. Purity of the isolated RNA was estimated by NanoDrop spectrophotometer (Thermo Scientific) and integrity was checked by running it on a 1% Tris-EDTA agarose gel.

Approximately 100 ng of total RNA was used per sample for cDNA synthesis. DNAse treatment and first strand synthesis were performed as described previously (Pathak et al., 2015). Quantitative real time PCRs (qPCRs) were performed in a total volume of 10 μl with Kapa SYBR Fast qPCR kit (KAPA Biosystems, Wilmington, MA) on an ABI 7500 fast machine operated with ABI 7500 software (Applied Biosystems). Technical duplicates were performed for each qPCR reaction. The fold change of gene expression in any experimental condition relative to wild-type was calculated as 2^−ΔΔ^^Ct^ where ΔΔCt = [Ct (target gene) − Ct (rp49)] Expt. − [Ct (target gene) − Ct (rp49)]. Primers specific for rp49 and ac5c were used as internal controls. Sequences of all primers used have been provided as Supplementary Table 2.

### Western blots

Larval CNS of appropriate genotypes were dissected in cold PBS. Between 5 and 10 brains were homogenized in 50 μl of NETN buffer [100mM NaCl, 20 mM Tris-Cl pH 8.0, 0.5 mM EDTA, 0.5% Triton-x-100, 1X Protease inhibitor cocktail (Roche) The homogenate (10–15 μl) was run on a 15% SDS-polyacrylamide gel. The protein was transferred to a 0.22 μm PVDF membrane by standard semi-dry transfer protocols (10V for 10 minutes). The membrane was incubated in the primary antibody overnight at 4°C. Primary antibodies were used at the following dilutions: Rabbit anti-H3K36me3 1:5000 (Abcam-ab9050). Rabbit anti-H3 1:5000 (Abcam-ab12079). Secondary antibodies conjugated with horseradish peroxidase were used at dilution of 1:3000 (anti-rabbit HRP; 32260, Thermo Scientific). Protein was detected by a chemiluminescent reaction (WesternBright ECL, Advansta K12045-20). Blots were first probed for H3K36me3, stripped with a solution of 3% glacial acetic acid for 10 mins, followed by re-probing with the anti-H3 antibody.

### Imaging

Third instar larval brains were dissected in insect hemolymph-like saline (HL3) (70 mM NaCl, 5 mM KCl, 20 mM MgCl_2_, 10 mM NaHCO_3_, 5 mM trehalose, 115 mM sucrose, 5 mM HEPES, 1.5 mM Ca2+, pH 7.2) after embedded in a drop of 0.1% low-melting agarose (Invitrogen). Embedded brains were bathed in HL3. GCaMP6m was used as the genetically encoded calcium sensor. Images were taken as a time series on an XY plane at an interval of 1 s using a 20x objective with an NA of 0.7 on an Olympus FV1000 inverted confocal microscope (Olympus Corp., Japan). 488 nm laser line was used to record GCaMP6m Ca^2+^ measurements. All live imaging experiments were performed with at least 10 independent brain preparations. For creating the effect of an acute loss of amino acid levels, we incubated the ex vivo preparations in either 0.5XEssential Amino Acids (EAA), obtained from a 50XEAA mixture lacking glutamine (Thermo Fisher Scientific) dissolved in HL3. At the point of withdrawal, the amino acid levels were diluted 10-fold using more HL3 thus creating the effect of amino acids withdrawal. For the dilution experiments, images were taken as a time series on an *xy* plane across 6 *z* planes at an interval of 4 s using a 10× objective with an NA of 0.4 and an optical zoom of 4 on an SP5 inverted confocal microscope mounted with a resonant scanner (Leica Microsystems). The *z* project across time was then obtained as a time series.

The raw images were extracted using FIJI (based on ImageJ version 2.1.0/1.53c; Schindelin et al., 2012). ΔF/F was calculated from selected regions of interest (ROI) using the formula ΔF/F = (F_t_-F_0_)/F_0_, where F_t_ is the fluorescence at time t and F_0_ is baseline fluorescence corresponding to the average fluorescence over the first 40-time frames. Mean ΔF/F time-lapses were plotted using MATLAB (R2019b; License number-1122786). A shaded error bar around the mean indicates the 95% confidence interval for CCh (50 μM) responses and the standard error of the mean (SEM) for amino acid withdrawal responses. Area under the curve was calculated from the point of stimulation which was considered as 0^th^ second for stimulation up to 300 s using Microsoft Excel and plotted using BoxPlotR (Spitzer et al., 2014).

### Immunohistochemistry

CNS were dissected in cold PBS, fixed with 4% PFA, washed with 0.2% PTX, blocked and incubated overnight in primary chick anti-GFP antibody (1:10,000; Abcam-Ab-13970) and rabbit anti-H3K36me3 1:5000 (Abcam-ab9050). They were then washed and incubated with anti-chick AlexaFluor-488 (#A1108, Invitrogen, RRID:AB_143165), anti-rabbit AlexaFluor-568 (#A1108, Invitrogen, RRID:AB_143165), (1:5000) DAPI, and mounted in 70% glycerol. Confocal images were obtained on the Confocal FV3000 microscope (Olympus) with a 40X, 0.7 NA objective. Images were visualized on FIJI (based on ImageJ version 2.1.0/1.53c)

### Cell sorting

Fluorescence activated cell sorting (FACS) was used to enrich cells from larval CNS, where neurons of interest were genetically labeled with GFP using the GAL4/UAS system (ref). The following genotypes were used for sorting: Wild type (*vGlut^VGN6341^GAL4>UAS-eGFP*); *itpr^IR^* (*vGlut^VGN6341^GAL4>itpr RNAi*); dSET2^IR-1^ (*vGlut^VGN6341^GAL4>dSET2IR-1); itpr^IR^; dSET2^OE^* (*vGlut^VGN6341^GAL4>itpr RNAi; UAS-dSET2*). Approximately 50 third instar larval CNSs per sample were washed in 1X PBS, 70% ethanol. Larval CNS were dissected in Schneider’s medium (Thermo Fisher Scientific) supplemented with 10% fetal bovine serum, 2% PenStrep, 0.02 mM insulin, 20 mM glutamine and 0.04 mg/mL glutathione. Post dissection, the larval CNS were treated with an enzyme solution (0.75 g/l collagenase and 0.4 g/l dispase in Rinaldini’s solution (8 mg/mL NaCl, 0.2 mg/mL KCl, 0.05 mg/mL NaH_2_PO_4_, 1 mg/mL NaHCO_3_, 0.1 mg/mL glucose) at room temperature for 30 min. They were then washed and resuspended in cold Schneider’s medium and gently triturated several times using a pipette tip to obtain a single-cell suspension. This suspension was then passed through a 40 μm mesh filter to remove clumps and kept on ice until sorting (less than an hour). Flow cytometry was performed on a FACS Aria Fusion cell sorter (BD Biosciences) with a 100 μm nozzle at 60 psi. The threshold for GFP-positive cells was set using dissociated neurons from a non GFP-expressing WT strain (Canton S). The same gating parameters were used to sort other genotypes in the experiment. GFP-positive cells were collected directly in Trizol and then frozen immediately in dry ice until further processing.

### RNA Sequencing and analysis of H3K36me3 marks

RNA-seq from larval CNSs: RNA isolation was performed from 15 larval CNSs of 84h AEL larvae of both control (*UAS-dSET2 IR/+)* and *dSET2 KD* (*vGlut^VGN6341GAL4^>UAS-dSET2^IR^*) genotypes using Trizol (Thermo Fisher Scientific) following the manufacturer’s protocol. Libraries with 500 ng total RNA per sample were prepared as described previously (Richhariya et al., 2017). Libraries were run on a Hiseq2500 platform. Biological triplicates were used for control and *dSET2 KD*. *itpr KD* RNA-seq data (GEO Accession number-GSE109637) were obtained from prior work (Jayakumar et al., 2018) and analyzed in the same manner. 50-70 million unpaired sequencing reads per sample were aligned to the dm3 release of the Drosophila genome using HISAT2 (Kim et al., 2015, 2019) and an overall alignment rate of 95.2-96.8% was obtained for all samples. Featurecounts (Liao et al., 2014) was used to assign the mapped sequence reads to the genome and obtain read counts. Differential expression analysis was performed using two independent methods: DESeq2 (Love et al., 2014) and edgeR (Robinson et al., 2009). A fold change cutoff of a minimum 2-fold change was used. Significance cutoff was set at an FDR-corrected p value of 0.05 for DESeq2 and edgeR.

Volcanoplots were generated using VolcaNoseR (Goedhart and Luijsterburg, 2020; https://huygens.science.uva.nl/). Comparison of gene lists and generation of Venn diagrams was performed using Whitehead BaRC public tools (http://jura.wi.mit.edu/bioc/tools/). Gene ontology analysis for molecular function was performed using DAVID (Huang et al., 2009b, 2009a). 277 downregulated genes common to *itpr KD* and *dSET2 KD* were used as the target set and all genes in Drosophila were used as background. Developmental gene expression levels were measured for downregulated genes using DGET (Hu et al., 2017) (www.flyrnai.org/tools/dget/web/) and plotted as a heatmap using ClustVis (Metsalu & Vilo, 2015) (https://biit.cs.ut.ee/clustvis/).

H3K36me3 enrichment data was obtained from a ChIP-chip dataset (ID_301) generated in *Drosophila* ML-DmBG3-c2 cells (Karpen et al., 2009) submitted to modEncode (Celniker et al., 2009). Enrichment scores for genomic regions were calculated using ‘computematrix’ and plotted as a tag density plot using ‘plotHeatmap’ from deeptools2 (Ramírez et al., 2016). All genes were scaled to 2kb and with a flanking region of 250 base pairs on either end. A 50 base pair length, of non-overlapping bins were used for averaging the score over each region length. Genes were sorted based on mean enrichment scores and displayed on the heatmap in descending order. For each downregulated gene, fold change upon *itpr KD, dSET2 KD* and relative enrichment H3K36me3 were compared and clustered using ClustVis (Metsalu & Vilo, 2015) (https://biit.cs.ut.ee/clustvis/). For each row of genes, row centering and unit variance scaling were applied prior to plotting. Genes were clustered based on Pearson’s correlation and ordered based on highest mean value. Two primary clusters emerged- High H3K36me3 with greater extent of downregulation and Low H3K36me3 with a lesser extent of downregulation.

### Phylogenetic Tree

Amino acid sequences for different H3K36 methyltransferases belonging to yeast, human and Drosophila species were obtained from UniprotKB (https://www.uniprot.org/). Multiple sequence alignment for the sequences was performed using the alignment tool ClustalW. A phylogenetic tree was constructed based on the neighbor joining method using ClustalW2 Phylogeny tool.

### Statistics

Datasets were evaluated for normality using the Kolmogorov-Smirnov test. Normally distributed datasets were compared using ANOVA followed with post hoc-Tukey test. Non-parametrically distributed datasets were evaluated using the Kruskall-Wallis test, followed with the Mann-Whitney U-test for pairwise comparisons. Pairwise comparisons for parametrically distributed datasets were performed using the 2-tailed Student’s t-test.

## Acknowledgements

We thank the Central Imaging and Flow Cytometry Facility (CIFF, NCBS) for maintenance and use of microscopes and the *Drosophila* facility (Flyfacility, NCBS) for stock maintenance and development of transgenics. We also thank members of the Notani Lab for discussions relating to ChIP-Seq analysis.

## Competing Interests

The authors declare no competing interests.

## Funding

This work was funded by grant No. BT/PR28450/MED/122/166/2018 from the Department of Biotechnology, Govt. of India and core support by NCBS, TIFR. RM and SR received graduate student fellowships from NCBS, TIFR and SJ was supported by a fellowship from CSIR, Govt. of India.

## Data availability

The RNA Sequencing data has been submitted to GEO (accession number-GSE162094) and is available here: https://www.ncbi.nlm.nih.gov/geo/query/acc.cgi?acc=GSE162094

## References

Bannister, A. J., Schneider, R., Myers, F. A., Thorne, A. W., Crane-Robinson, C., & Kouzarides, T. (2005). Spatial distribution of di-and tri-methyl lysine 36 of histone H3 at active genes. Journal of Biological Chemistry 280, 17732–17736. https://doi.org/10.1074/jbc.M500796200

Boulan, L., Milán, M., & Léopold, P. (2015). The systemic control of growth. Cold Spring Harbor Perspectives in Biology 7, 1–30. https://doi.org/10.1101/cshperspect.a019117

Britton, J. S., Lockwood, W. K., Li, L., Cohen, S. M., & Edgar, B. A. (2002). Drosophila’s insulin/PI3-kinase pathway coordinates cellular metabolism with nutritional conditions. Developmental Cell 2, 239–249. https://doi.org/10.1016/S1534-5807(02)00117-X

Celniker, S. E., Dillon, L. A. L., Gerstein, M. B., Gunsalus, K. C., Henikoff, S., Karpen, G. H., Kellis, M., Lai, E. C., Lieb, J. D., MacAlpine, D. M., Micklem, G., Piano, F., Snyder, M., Stein, L., White, K. P., & Waterston, R. H. (2009). Unlocking the secrets of the genome. Nature 459, 927–930. https://doi.org/10.1038/459927a

Chen, T., Wardill, T. J., Sun, Y., Pulver, S. R., Renninger, S. L., Baohan, A., Schreiter, E. R., Kerr, R. A., Orger, M. B., Jayaraman, V., Looger, L. L., Svoboda, K., & Kim, D. S. (2013). Ultrasensitive fluorescent proteins for imaging neuronal activity. Nature 499, 295–300. https://10.1038/nature12354

Chen, X., Rahman, R., Guo, F., & Rosbash, M. (2016). Genome-wide identification of neuronal activity-regulated genes in *Drosophila*. Elife 5, e19942 https://doi.org/10.7554/eLife.19942

Dolmetsch, R. (2003). Excitation-transcription coupling: signaling by ion channels to the nucleus. Science’s STKE 2003, pe4. https://doi:10.1126/stke.2003.166.pe4

Egorova, P. A., & Bezprozvanny, I. B. (2018). Inositol 1,4,5-trisphosphate receptors and neurodegenerative disorders. FEBS Journal 285, 3547–3565. https://doi.org/10.1111/febs.14366

Goedhart, J., Luijsterburg, M.S. (2020). VolcaNoseR is a web app for creating, exploring, labeling and sharing volcano plots. Scientific Reports 10, 20560 https://doi.org/10.1038/s41598-020-76603-3

Hasan, G., & Sharma, A. (2020). Regulation of neuronal physiology by Ca2+release through the IP3R. Current Opinion in Physiology 17, 1–8. https://doi.org/10.1016/j.cophys.2020.06.001

Hu, Y., Comjean, A., Perrimon, N., & Mohr, S. E. (2017). The Drosophila Gene Expression Tool (DGET) for expression analyses. BMC Bioinformatics 18, 1–9. https://doi.org/10.1186/s12859-017-1509-z

Huang, D. W., Sherman, B. T., & Lempicki, R. A. (2009a). Bioinformatics enrichment tools: Paths toward the comprehensive functional analysis of large gene lists. Nucleic Acids Research 37, 1–13. https://doi.org/10.1093/nar/gkn923

Huang, D. W., Sherman, B. T., & Lempicki, R. A. (2009b). Systematic and integrative analysis of large gene lists using DAVID bioinformatics resources. Nature Protocols 4, 44–57. https://doi.org/10.1038/nprot.2008.211

Iossifov, I., O’Roak, B. J., Sanders, S. J., Ronemus, M., Krumm, N., Levy, D., Stessman, H. A., Witherspoon, K. T., Vives, L., Patterson, K. E., Smith, J. D., Paeper, B., Nickerson, D. A., Dea, J., Dong, S., Gonzalez, L. E., Mandell, J. D., Mane, S. M., Murtha, M. T.,… Wigler, M. (2014). The contribution of de novo coding mutations to autism spectrum disorder. Nature 515, 216–221. https://doi.org/10.1038/nature13908

Jayakumar, S., Richhariya, S., Deb, B. K., & Hasan, G. (2018). A multicomponent neuronal response encodes the larval decision to pupariate upon amino acid starvation. Journal of Neuroscience 38, 10202–10219. https://doi.org/10.1523/JNEUROSCI.1163-18.2018

Jayakumar, S., Richhariya, S., Reddy, O. V., Texada, M. J., & Hasan, G. (2016). Drosophila larval to pupal switch under nutrient stress requires IP 3R / Ca2+ signalling in glutamatergic interneurons. eLife 5, e17495 https://doi.org/10.7554/eLife.17495

Kharchenko, P. V., Alekseyenko, A. A., Schwartz, Y. B., Minoda, A., Riddle, N., Ernst, J., Sabo, P. J., Larschan, E., Gorchakov, A. A., Gu, T., Linder-Basso, D., Plachetka, A., Shanower, G., Tolstorukov, M. Y., Luquette, L. J., Xi, R., Jung, Y. L., Park, R. W., Bishop, E. P.,… Park, P. J. (2011). Comprehensive analysis of the chromatin landscape in Drosophila melanogaster. Nature 471, 480–486. https://doi.org/10.1038/nature09725

Kim, D., Langmead, B., & Salzberg, S. L. (2015). HISAT: A fast spliced aligner with low memory requirements. Nature Methods 12, 357–360. https://doi.org/10.1038/nmeth.3317

Kim, D., Paggi, J. M., Park, C., Bennett, C., & Salzberg, S. L. (2019). Graph-based genome alignment and genotyping with HISAT2 and HISAT-genotype. Nature Biotechnology 37, 907–915. https://doi.org/10.1038/s41587-019-0201-4

Kizer, K. O., Phatnani, H. P., Shibata, Y., Hall, H., Greenleaf, A. L., & Strahl, B. (2005). A Novel Domain in Set2 Mediates RNA Polymerase II Interaction and Couples Histone H3 K36 Methylation with Transcript Elongation. Molecular and cellular biology 25, 3305–3316. https://doi.org/10.1128/MCB.25.8.3305-3316.2005

Kouzarides, T. (2007). Chromatin Modifications and Their Function. Cell 128, 693–705. https://doi.org/10.1016/j.cell.2007.02.005

Liao, Y., Smyth, G. K., & Shi, W. (2014). FeatureCounts: An efficient general purpose program for assigning sequence reads to genomic features. Bioinformatics 30, 923–930. https://doi.org/10.1093/bioinformatics/btt656

Love, M. I., Huber, W., & Anders, S. (2014). Moderated estimation of fold change and dispersion for RNA-seq data with DESeq2. Genome Biology 15, 1–21. https://doi.org/10.1186/s13059-014-0550-8

Lumish, H. S., Wynn, J., Devinsky, O., & Chung, W. K. (2015). Brief Report: SETD2 Mutation in a Child with Autism, Intellectual Disabilities and Epilepsy. Journal of Autism and Developmental Disorders 45, 3764–3770. https://doi.org/10.1007/s10803-015-2484-8

McDaniel, S. L., Hepperla, A. J., Huang, J., Dronamraju, R., Adams, A. T., Kulkarni, V. G., Davis, I. J., & Strahl, B. D. (2017). H3K36 Methylation Regulates Nutrient Stress Response in Saccharomyces cerevisiae by Enforcing Transcriptional Fidelity. Cell Reports 19, 2371–2382. https://doi.org/10.1016/j.celrep.2017.05.057

Metsalu, T., & Vilo, J. (2015). ClustVis: A web tool for visualizing clustering of multivariate data using Principal Component Analysis and heatmap. Nucleic Acids Research 43, W566–W570. https://doi.org/10.1093/nar/gkv468

Mirth, C., Truman, J. W., & Riddiford, L. M. (2005). The role of the prothoracic gland in determining critical weight for metamorphosis in Drosophila melanogaster. Current Biology 15, 1796–1807. https://doi.org/10.1016/j.cub.2005.09.017

Nijhout, H. F. (2003). The control of body size in insects. Developmental Biology, 261, 1–9. https://doi.org/10.1016/S0012-1606(03)00276-8

Passani, L. A., Bedford, M. T., Faber, P. W., McGinnis, K. M., Sharp, A. H., Gusella, J. F., Vonsattel, J. P., & MacDonald, M. E. (2000). Huntingtin’s WW domain partners in Huntington’s disease post-mortem brain fulfill genetic criteria for direct involvement in Huntington’s disease pathogenesis. Human Molecular Genetics 9, 2175–2182. https://doi.org/10.1093/hmg/9.14.2175

Ramírez, F., Ryan, D. P., Grüning, B., Bhardwaj, V., Kilpert, F., Richter, A. S., Heyne, S., Dündar, F., & Manke, T. (2016). deepTools2: a next generation web server for deep-sequencing data analysis. Nucleic Acids Research 44, W160–W165. https://doi.org/10.1093/nar/gkw257

Richhariya, S., Jayakumar, S., Abruzzi, K., Rosbash, M., & Hasan, G. (2017). A pupal transcriptomic screen identifies Ral as a target of store-operated calcium entry in Drosophila neurons. Scientific Reports 7, 1–12. https://doi.org/10.1038/srep42586

Robinson, M. D., McCarthy, D. J., & Smyth, G. K. (2009). edgeR: A Bioconductor package for differential expression analysis of digital gene expression data. Bioinformatics 26, 139–140. https://doi.org/10.1093/bioinformatics/btp616

Schaft, D., Roguev, A., Kotovic, K. M., Shevchenko, A., Sarov, M., Shevchenko, A., Neugebauer, K. M., & Stewart, A. F. (2003). The histone 3 lysine methyltransferase, SET2, is involved in transcriptional elongation. Nucleic Acids Research 31, 2475–2482. https://doi.org/10.1093/nar/gkg372

Schindelin, J., Arganda-Carreras, I., Frise, E., Kaynig, V., Longair, M., Pietzsch, T., Preibisch, S., Rueden, C., Saalfeld, S., Schmid, B., Tinevez, J. Y., White, D. J., Hartenstein, V., Eliceiri, K., Tomancak, P., & Cardona, A. (2012). Fiji: An open-source platform for biological-image analysis. Nature Methods 9, 676–682. https://doi.org/10.1038/nmeth.2019

Sharma, A., Nguyen, H., Geng, C., Hinman, M. N., Luo, G., & Lou, H. (2014). Calcium-mediated histone modifications regulate alternative splicing in cardiomyocytes. Proceedings of the National Academy of Sciences of the United States of America 111, E4920–E4928. https://doi.org/10.1073/pnas.1408964111

Stabell, M., Larsson, J., Aalen, R. B., & Lambertsson, A. (2007). *Drosophila* dSet2 functions in H3-K36 methylation and is required for development. Biochemical and Biophysical Research Communications 359, 784–789. https://doi.org/10.1016/j.bbrc.2007.05.189

Wang, C. M., Tsai, S. N., Yew, T. W., Kwan, Y. W., & Ngai, S. M. (2010). Identification of histone methylation multiplicities patterns in the brain of senescence-accelerated prone mouse 8. Biogerontology 11, 87–102. https://doi.org/10.1007/s10522-009-9231-5

Wang, Y. P., & Lei, Q. Y. (2018). Metabolite sensing and signaling in cell metabolism. Signal Transduction and Targeted Therapy 3, 1–9. https://doi.org/10.1038/s41392-018-0024-7

Ward, P. S., & Thompson, C. B. (2012). Signaling in control of cell growth and metabolism. Cold Spring Harbor Perspectives in Biology 4, 1–15. https://doi.org/10.1101/cshperspect.a006783

Whitlock, J. P., Galeazzi, D., & Schulman, H. (1983). Acetylation and calcium-dependent phosphorylation of histone H3 in nuclei from butyrate-treated HeLa cells. Journal of Biological Chemistry 258, 1299–1304.

